# Robust analysis of prokaryotic pangenome gene gain and loss rates with Panstripe

**DOI:** 10.1101/2022.04.23.489244

**Authors:** Gerry Tonkin-Hill, Rebecca A Gladstone, Anna K Pöntinen, Sergio Arredondo-Alonso, Stephen D Bentley, Jukka Corander

## Abstract

Horizontal gene transfer (HGT) plays a critical role in the evolution and diversification of many microbial species. The resulting dynamics of gene gain and loss can have important implications for the development of antibiotic resistance and the design of vaccine and drug interventions. Methods for the analysis of gene presence/absence patterns typically do not account for errors introduced in the automated annotation and clustering of gene sequences. In particular, methods adapted from ecological studies, including the pangenome gene accumulation curve, can be misleading as they may reflect the underlying diversity in the temporal sampling of genomes rather than a difference in the dynamics of HGT. Here, we introduce Panstripe, a method based on Generalised Linear Regression that is robust to population structure, sampling bias and errors in the predicted presence/absence of genes. We demonstrate using simulations that Panstripe can effectively identify differences in the rate and number of genes involved in HGT events, and illustrate its capability by analysing several diverse bacterial genome datasets representing major human pathogens. Panstripe is freely available as an R package at https://github.com/gtonkinhill/panstripe.

## Introduction

Genetic variation within microbial populations is shaped by both the accumulation of variation from point mutations as well as by the acquisition and loss of genetic material through Horizontal Gene Transfer (HGT). HGT can occur via the uptake of DNA from the environment, with the help of Mobile Genetic Elements (MGEs) (phages, integrative conjugative elements and plasmids) or from direct contact between bacterial cells (1). Genes are also frequently duplicated and lost vertically upon cell division (2). The influence of these sources of variation varies dramatically by species. Clonal species such as *Mycobacterium tuberculosis* typically accumulate variation nearly entirely through point mutations while naturally transformable species such as *Streptococcus pneumoniae* and *Neisseria meningitidis* have very high rates of homologous recombination (3). In other species such as *Salmonella enterica*, horizontal exchange is generally restricted to the movement of MGEs (4). While HGT does not always have an impact on a microbe’s fitness, it can lead to critical phenotypic changes such as the acquisition of antimicrobial resistance, virulence factors and vaccine escape (5, 6). A common approach to analysing horizontal exchange in microbial genomics is to group homologous gene sequences into orthologous and paralogous gene clusters. The union of these clusters within a particular species or group is commonly referred to as the pangenome (7). Genes are often further classified into either the ‘core’ genome which are found in almost all members of the group and the ‘accessory’ genome which are only found in a subset of genomes. Species with a limited accessory genome such that all genes are likely to have already been observed are often described as ‘closed’ while species with a diverse accessory genome such that each additional genome contributes novel genes are described as ‘open’.

A number of tools have been developed to infer a pangenome given a collection of annotated genomes (8–13). A common output of these approaches is a binary gene presence/absence matrix where genomes are represented by rows and orthologous gene clusters by columns. After generating a gene presence/absence matrix, researchers are often interested in comparing the size of pangenomes between datasets, determining the rate of horizontal gene exchange as well as identifying whether a pangenome is ‘open’ or ‘closed’.

To answer these questions, researchers frequently generate a gene accumulation curve as is often done in ecological studies of species diversity (7, 14). Here, the number of unique gene clusters identified is plotted against the number of genomes. The order in which the genomes are considered is usually permuted randomly to account for the variation that this order could have on the resulting plot. In some cases, a power law such as Heaps’ or Zipf’s law is fit to this curve to give a parameter estimate of the diminishing number of new genes found with each additional genome and to determine whether the pangenome is open or closed (15).

A neglected problem with this approach is that it fails to account for the underlying diversity of the set of sampled genomes. For example, a set of genomes taken from within an outbreak is likely to involve far fewer gene exchange events than a diverse sample from a species with thousands of years of evolution separating isolates. Methods that make use of a phylogeny constructed from the genetic diversity present in genes found in all the genomes (the ‘core’ genome) help to address this issue by controlling for the underlying diversity of the sample. The branch lengths of the core genome phylogeny indicate the evolutionary time over which gene gain and loss events could have occurred. Shorter branch lengths separating more closely related taxa would be expected to have fewer associated gene exchange events. Methods that rely on construction of such a phylogeny include those based on, maximum parsimony (16), maximum likelihood (17–19) and Bayesian phylogenetics (20). Two notable models that use this approach are the Infinitely Many Genes (IMG) model and the Finitely Many Genes model (FMG) (21–24). The IMG model assumes an infinite pool of genes and that a particular gene can only be gained once while the FMG model assumes that genes belong to a finite pool and that multiple gene gain and loss events of the same gene can occur. Many models also collapse paralogous clusters into gene families before the inference of gene gain and loss rates (16, 17, 19). A significant limitation of these approaches is that they generally assume that there is no error in the inferred pangenome presence/absence matrix. We and others have shown that gene annotation errors and the complexities of clustering genes into orthologous families can introduce substantial numbers of erroneous gene clusters (12, 13, 19, 25). While a subset of models do account for errors in the predicted presence/absence of genes, these have mostly been optimised for the analysis of eukaryotes and focus on a small number of gene families involving multiple genes (19). Most models also make the simplifying assumption that genes are gained or lost individually which can significantly bias estimates of the rate of gene exchange, particularly when the exchange of MGEs is frequent.

To address these limitations we have developed Panstripe, an approach that compares the rates of core and accessory genome evolution to account for both population structure and errors in the pangenome gene presence/absence matrix. We demonstrate using extensive simulations, and by analysing a diverse range of bacterial genome datasets that Panstripe is: (i) able to effectively compare the gene exchange rates between pangenomes, (ii) can be used to determine if there is a temporal signal in the accessory genome, and (iii) can identify if the size of gene exchange events differs between pangenomes.

## Results

### Overview

Panstripe accepts a phylogeny produced using standard pipelines and a corresponding gene presence/absence matrix as produced by most pangenome inference tools (10, 12, 26). The length of each branch in the phylogeny is then compared to the number of gene gain and loss events inferred to have occurred on that branch using a Generalised Linear Model (GLM) (Figure 1). The ancestral gene gain and loss events on each branch can be inferred using common Ancestral State Reconstruction (ASR) methods including maximum parsimony, maximum likelihood and stochastic mapping (17, 27–29). While this is a critical step in the Panstripe algorithm, we have found that the approach is robust to the choice of ASR method (Results). The Panstripe algorithm is similar to root-to-tip regression, which is used to test for temporal signal in phylogenies (30). However, in contrast to TempEst, the regression is not performed on the root-to-tip distance but rather on the individual branch lengths. This avoids the problematic dependence structure which makes the root-to-tip regression not suitable for statistical hypothesis testing (30, 31).

**Fig. 1.**
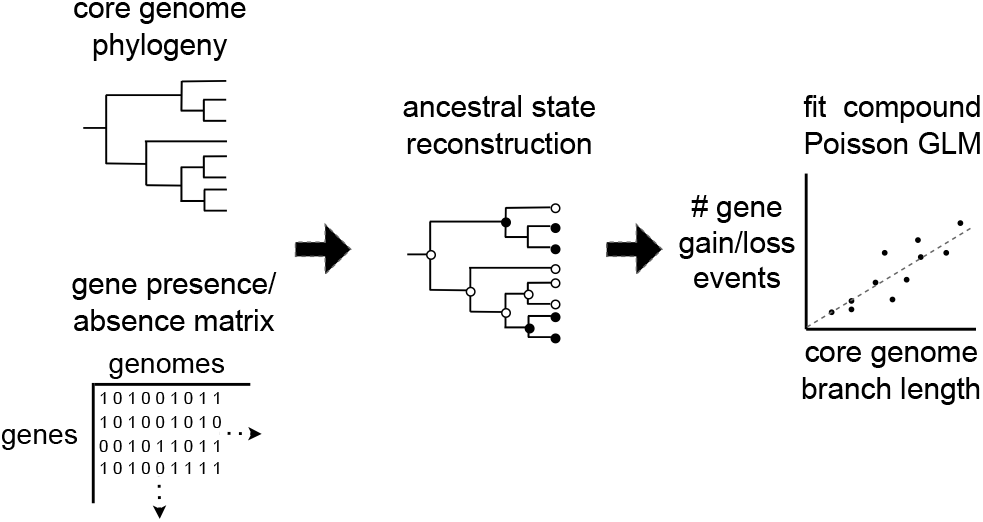
A schematic of the Panstripe algorithm. A binary gene presence/absence matrix and a core genome phylogeny are taken as input. The ancestral states of each gene are then determined before fitting a compound Poisson GLM to compare the core branch length with the number of gene gain and loss events on each branch. Additional terms are included to account for the depth of a branch and whether or not it occurs at the tips of the phylogeny.

Panstripe assumes that the number of gene gain and loss events on each branch are distributed according to a Compound Poisson distribution. This allows for multiple genes to be gained and lost in a single event (see Methods). Errors in the gene presence/absence matrix introduced at either the sequencing, assembly, annotation or pangenome clustering stage are unlikely to correlate with the core genome phylogeny. That is, we expect to see a similar number of errors on shorter terminal branches originating from a more closely related clade as a long terminal branch leading to a taxon with no close relatives. Ancestral state reconstruction of these errors will place them at the terminal branches (tips) of the phylogeny. By including a binary covariate which indicates whether or not a branch is located at the tip of the phylogeny we can control for the presence of errors in the pangenome. Without careful consideration of the individual gene sequences, it is not possible to accurately distinguish errors from very rare genes that are observed once. As a result, Panstripe groups the signal from both these sources into a single term.

The GLM framework used by Panstripe can also be used to compare the rates of HGT between pangenomes or to identify associations with additional covariates of interest. This relies on the given phylogenies having the same scale which can be achieved using time-scaled trees. Alternatively, SNP-scaled phylogenies can be used if differences in the ratio of core to accessory genome variation are of interest. To test whether the number of genes involved in each recombination event is significantly different between two pangeomes, Panstripe allows different models to be fitted separately to the mean and dispersion structure of the GLM (32).

The Panstripe package provides several plotting functions including an alternative to the popular pangenome accumulation curve that controls for both errors and differences in the core genome diversity of the underlying isolates. The package is written in R and is available freely under an MIT licence from https://github.com/gtonkinhill/panstripe.

### Panstripe is robust to errors in pangenome clustering

To assess the robustness of the Panstripe algorithm to the choice of pangenome clustering method, we considered the pangenome analysis of a large outbreak of highly clonal *Mycobacterium tuberculosis* (Mtb) in London spanning 14 years (33). This collection of genomes has been previously analysed using both the Roary and Panaroo pangenome clustering pipelines (12). Due to the very low mutation rate and highly clonal nature of the outbreak, we would expect there to be no pangenome variation in this dataset making it a useful control for assessing whether pangenome inference tools can account for errors.

Figure 2A presents the pangenome accumulation curves of the resulting gene presence/absence matrices output by the Roary and Panaroo pipelines. The very large difference in the two curves demonstrates that the accumulation curve method is highly sensitive to the different error rates of the two pipelines. We have previously shown that Panaroo can significantly reduce the number of errors when generating a pangenome clustering as observed in Figure 2A. However, Panaroo still estimates a small amount of accessory variation in this clonal Mtb dataset.

**Fig. 2.**
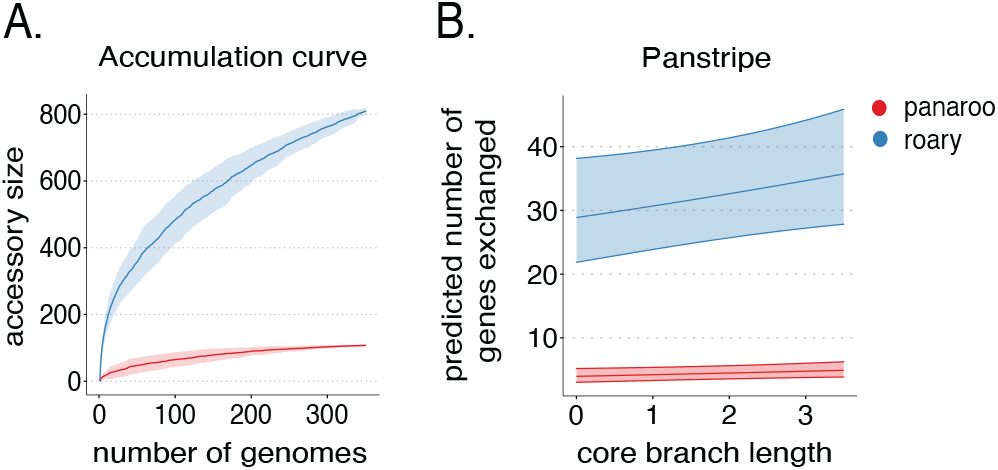
**A.)** Pangenome accumulation curves after running both Roary (blue) and Panaroo (red) on 351 Mtb genomes from an outbreak in London. The ribbon indicates the variation in the curve found by permuting the genome order 100 times. **B.)** Plot of the core branch length versus the predicted number of accessory genes for the same set of Mtb genomes according to the Panstripe model. The ribbon displays the 95% confidence interval of the Panstripe model fit and indicates that the inferred slopes are not significantly different from zero for both the Panaroo and Roary presence-absence matrices.

Figure 2B indicates that the Panstripe algorithm can accurately account for the differences in error rates of the two pipelines. By plotting the core branch lengths against the predicted number of gene gain and loss events it becomes clear that the large number of accessory genes identified by the Roary pipeline do not correlate with the core phylogeny and thus are likely to be erroneous. Panstripe correctly estimates a very similar association between the core genome branch length and the number of inferred gene gain and loss events on each branch for both the Roary and Panaroo pangenomes (*β*_*core*_ = 0.0621 and 0.0625 respectively). In both cases, Panstripe found that the coefficients were not significantly different from zero (p=0.871), which is consistent with the closed pangenome of Mtb. Panstripe can also be used to compare the differences in the rate of gene exchange events associated with the tips of the phylogeny. Here, it correctly identifies a significantly elevated rate of errors in the Roary gene presence/absence matrix (p<0.001).

### Panstripe outperforms alternative methods on simulated data

Simulations indicate that Panstripe is more robust to the impacts of errors and population structure than alternative methods including those based on information theory (15, 34) and phylogenetically informed approaches (22–24) (Figure 3, Supplementary Figure 3). To simulate pangenome datasets, we first generate core genome phylogenies before simulating the number and size of gene gain and loss events along each branch (see Methods). In line with the assumptions of Dollo’s parsimony, we assume an elevated frequency of gene loss relative to gene gain as such an asymmetry is frequently observed in bacterial genomes (35, 36). Errors in the pangenome matrix were generated by randomly adding or removing single entries in the matrix.

**Fig. 3.**
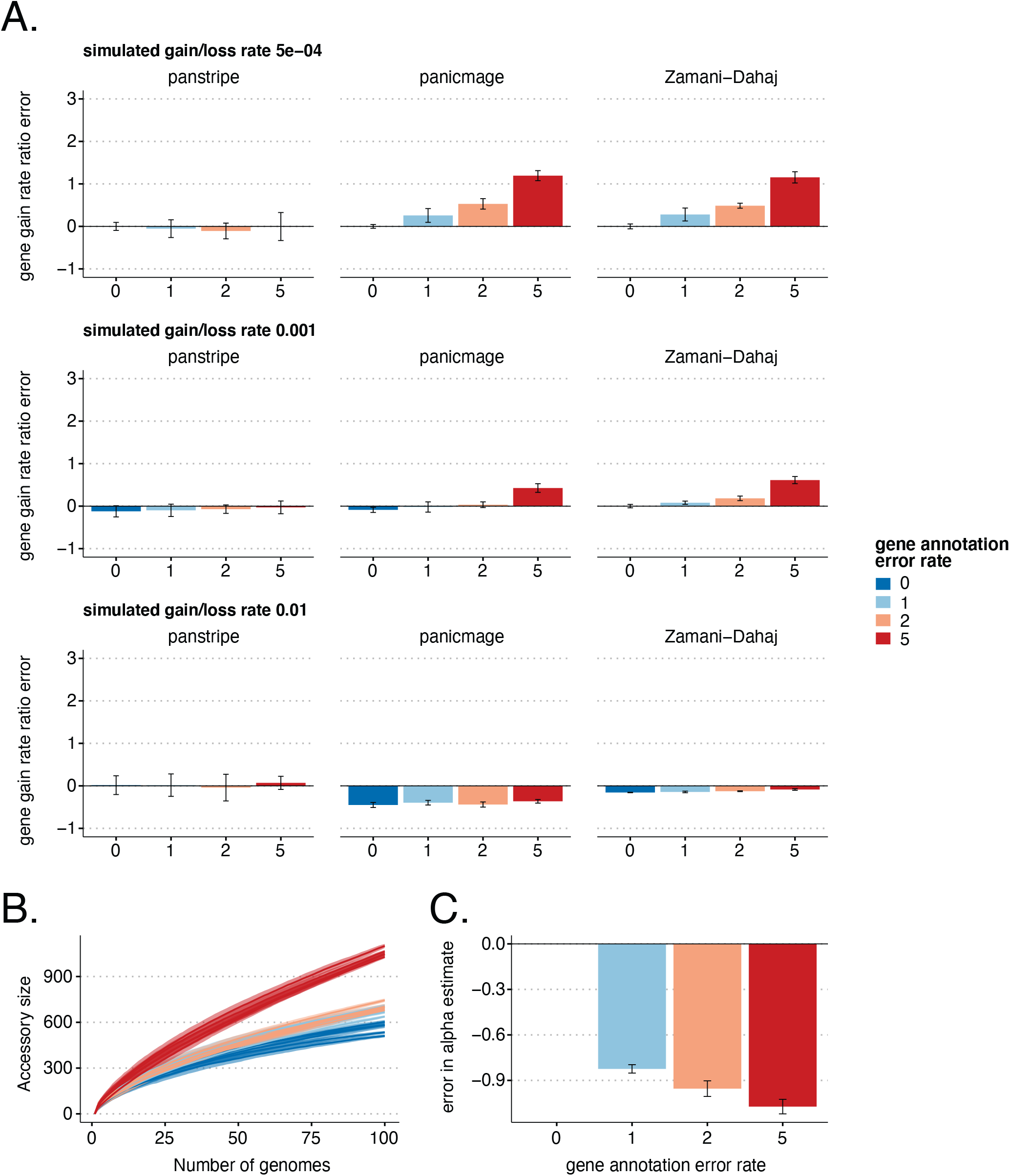
**A.)** Bars indicate the mean percentage error in the estimated ratio of gene gain and loss rate compared to the reference rate of 5 × 10^*−*4^ with a simulated annotation error of 0. The three rows represent increasingly large gene gain and loss rates of 5 × 10^*−*4^ 0.001 and 0.01. The simulated annotation error rates are given along the x-axis. **B.)** Pangenome accumulation curves with a simulated gene gain and loss rate of 0.001. The colours represent the increasing annotation error rates. **C.)** The corresponding error in the *α* parameter estimates after fitting Heaps law to the curves in B. Lower *α* estimates indicate a more ‘open’ pangenome. Thus, higher rates of annotation error can lead to incorrect estimates of whether a pangenome is open or closed.

It is challenging to compare the output of the various pangenome dynamics inference tools as each makes different assumptions and they do not attempt to infer the same set of parameters. To cope with this, we focused on the ability of each tool to accurately quantify differences in the dynamics of gene gain and loss between pangenome datasets. For those tools that infer a gene gain rate, we tested whether the ratio of inferred gene gain rates matched that of the simulations. Thus, we would expect a ratio of the estimated gene gain rate of 1 for a pair of simulations with the same parameters and a ratio of 2 if one simulation used a gene gain rate that was twice as large. This allows us to investigate the ability of each tool to distinguish the dynamics of gene gain and loss between datasets. Approaches based on information theory, such as the pangenome accumulation curve and fitting a Heaps power law do not have a parameter that is easily compared with the gene gain rate. Instead, we compare the results of these methods using a fixed gene gain and loss rate subject to different levels of error and bias.

Figure 3A indicates that Panstripe was the only tool that provided consistent parameter estimates both at higher error rates and across a range of gene gain and loss rates. The phylogenetic informed methods such as the IMG model of Panicmage and the FMG model of Zamani-Dahaj et al., both performed poorly at higher error rates. Interestingly, although these methods were generally accurate in the absence of errors in the presence-absence matrix, they both systematically underestimated higher gene gain rates. In our analyses, we could not get the IMG model of Collins et al., to give consistent results (Supplementary Figure 3). The information theory-based approaches were highly sensitive to errors in the presence-absence matrix (Figure 3B-C). Increased error rates led to both substantial differences in the slopes of the pangenome accumulation curve and an underestimate in the *α* parameter of Heaps power law. This suggests that under realistic error rates, the use of Heaps power to classify pangenomes into ‘open’ and ‘closed’ is problematic. To test whether these results held for different ratios of gene gain and loss we repeated the analysis assuming equal frequencies of gene gain and loss as well as an elevated rate of gene gain. In both cases we observed very similar results (Supplementary Figures 8 and 9).

Ancestral state reconstruction forms a critical component of the Panstripe algorithm. To test the sensitivity of the approach to the choice of algorithm, we simulated multiple pangenome datasets with increasing rates of annotation error. We then ran Panstripe using the included ASR algorithms (maximum parsimony, maximum likelihood and stochastic mapping). We found that Panstripe performed similarly using all three of the included algorithms with the variation in the estimated rate of gene gain and loss using the same set of simulation parameters exceeding the variation between the different ASR algorithms (Supplementary Figure 4A). These simulations represent relatively well-behaved datasets. When we compared the gene gain and loss estimates on the more challenging clonal Mtb dataset there was a larger difference between the different ASR algorithms and only maximum parsimony correctly indicated a gene gain and loss rate that was indistinguishable from zero (Supplementary Figure 4B-C). The very low temporal signal in the Mtb phylogeny leads to very short branch lengths and multichotomies. This causes the maximum likelihood and stochastic mapping algorithms to incorrectly assign gene gain and loss events to longer branches. Consequently, as the major goal of the Panstripe algorithm is to enable robust inference of gene gain and loss rates we make use of maximum parsimony by default. However, the program includes the option to use both maximum likelihood and stochastic mapping algorithms so that users can easily test the sensitivity of their analyses to the assumptions of these different algorithms.

Similar to the impacts of errors in the pangenome gene presence-absence matrix, we found that Panstripe was more robust to sampling biases and the underlying phylogenetic structure of a pangenome dataset. To test this, we simulated five large phylogenies and accompanying gene presenceabsence matrices. We then selected a small sub-clade (≥30 genomes) within each simulation to create a smaller, more closely related dataset. Here, the simulation parameters of the large dataset and the sub-clade are the same and thus the ratio of the estimated parameter for each method on each dataset should be 1. Figure 4A indicates that both panstripe and the phylogenetically informed method of Zamani-Dahaj et al., provided consistent parameter estimates of the full dataset and sub-clade. Conversely, the ratio of the gene gain rate estimated by Panicmage and the Heaps power law *α* parameter between the full dataset and the sub-clade both had error rates above 50%. Similarly, the pangenome accumulation curves (Figure 4B) were highly sensitive to sampling bias. Overall, these simulations indicate that panstripe outperforms other methods by providing gene gain and loss rate estimates that better reflect the true difference between datasets whilst being robust to both sampling bias and error in the gene presence-absence matrix.

**Fig. 4.**
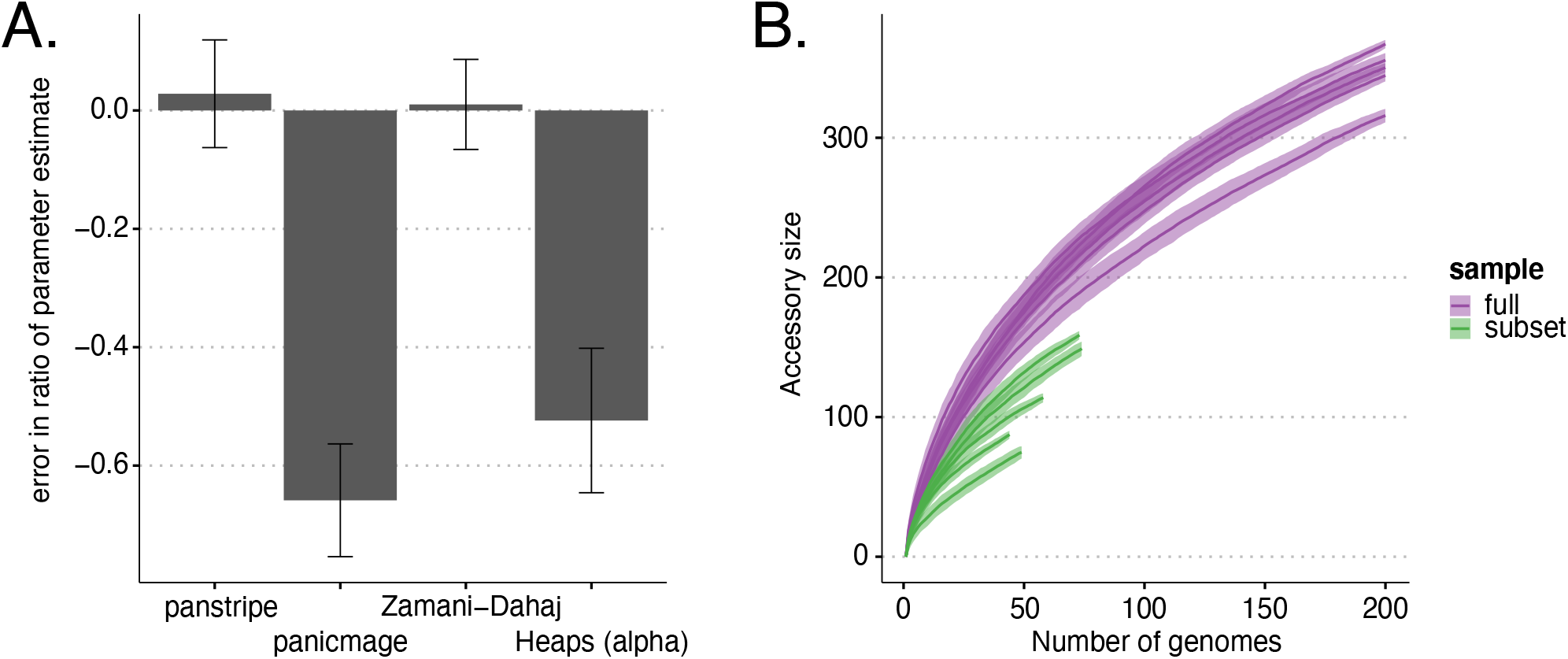
**A.)** Bars indicate the mean percentage error in the estimated ratio of gene gain and loss rate in a smaller sub-clade of the simulated data when compared to estimating the rate on the full simulation. Only Panstripe and the method of Zamanj-Dahaj et al., accurately reported similar parameters in both the small and large sets. This suggests that Panicmage and fitting a Heaps power law are highly sensitive to sampling biases in the data. **B.)** Similar to the Heaps power law, pangenome accumulation curves provide misleading differences when comparing the sub-clade and full datasets.

To verify that Panstripe can differentiate between different rates of annotation error we generated simulated pangenomes using three different gene gain and loss rates and five annotation error rates. Supplementary Figure 1 indicates the resulting p-values associated with the tip parameter of the Panstripe GLM. Panstripe was able to accurately differentiate different rates of annotation error for all datasets except those generated using the highest gene gain and loss rate. At the highest gene gain and loss rate, the number of real gene exchange events tends to dominate the impact of errors and thus it is more challenging for the Panstripe algorithm to distinguish different rates of annotation error. We also considered whether Panstripe can distinguish between the size of gene exchange events by simulating two gene gain and loss rates and four different recombination sizes. Supplementary Figure 2 shows that Panstripe was able to accurately differentiate between all four simulated recombination sizes.

### Panstripe accurately identifies within and between species differences in gene gain and loss rates

To compare the estimates of the Panstripe algorithm on species that are known to have a diverse accessory genome, we considered two previously described datasets. The first included 315 *Enterococcus faecalis* genomes from three major hospital-associated clades sampled in the Netherlands, Spain and Portugal (37). *E. faecalis* is both a commensal and nosocomial pathogen and can successfully inhabit a wide range of host niches. The generalist ecological lifestyle of the microbe is facilitated by a diverse accessory genome with little association between specific niches and particular accessory genes (38, 39). Instead, adaptation to the hospital-associated niche is thought to be due to selection for survivability in a broader set of niches (37). We generated pangenome curves and ran the Panstripe algorithm on three of the major hospital-associated *E. faecalis* clades pp18, pp2 and pp6 (Figure 5A). Although clade pp18 (including sequence types ST159 and ST525) appeared to have a larger accessory genome according to the pangenome accumulation curve (Figure 5B), the Panstripe algorithm revealed that this was likely to be driven by sampling biases as clade pp18 included a more diverse set of genomes (5C-D). Instead, the pp6 clade (mainly of ST28) had a higher rate of gene exchange when compared to both the pp2 (mainly of ST6) (p<0.001) and pp18 (p<0.001) clades. The pp6 clade is the most recent hospital-associated clade to have emerged in this dataset and thus the elevated rate of gene exchange could be a result of the additional selection pressures acting within the hospital environment which may drive the acquisition of antimicrobial resistance plasmids (37). Consistent with this hypothesis, panstripe found that the size of gene gain and loss events was higher in clade pp6.

**Fig. 5.**
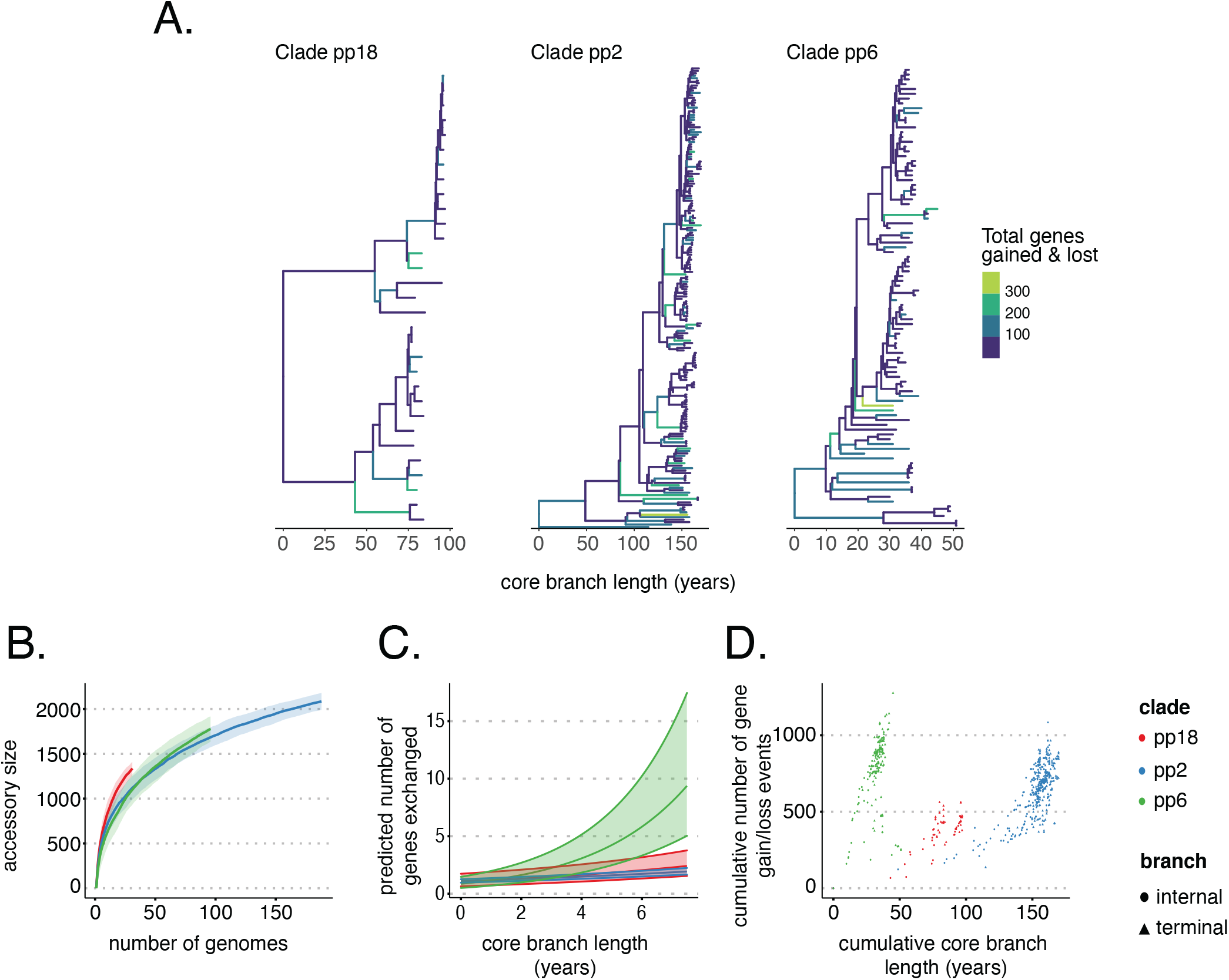
**A.)** Phylogenies of three major *E. faecalis* clades taken from Pöntinen et al., with branches coloured by the number of gene gain and loss events inferred using maximum parsimony. **B.)** The pangenome accumulation curves of the same clades using the pangenomes inferred using Panaroo in Pöntinen et al. **C.)** The predicted slope of the relationship between core genome branch length and the number of gene gain and loss events is inferred by the Panstripe algorithm. **D.)** The cumulative number of gene gain and loss events versus the cumulative branch length starting from the root node of each tree in A. This is a similar plot to the common ‘root-to-tip’ plot used in phylogenetic dating.

The second dataset consisted of four subclades (A, B, C1, C2) of the globally disseminated ST131 clone from a longitudinal study of *Escherichia coli* isolates from the Norwegian surveillance on resistant microbes (NORM) program (40). Similar to our analysis of *E. faecalis* the estimated pangenome accumulation curve for these clades gave strikingly different results to that produced by Panstripe (Figure 6A-C). According to the pangenome accumulation curve, Clade B exhibited the largest accessory genome diversity. However, this is likely due to the increased age of Clade B as its expansion in the Norwegian population occurred nearly a decade earlier than the other clades (40). Thus, the pangenome accumulation curve is again reflecting the underlying population structure of the dataset rather than a difference in the accessory genome dynamics of the clades. Instead, Panstripe predicts an increased rate of gene gain and loss for the C1 and C2 clades which expanded more recently and have a higher prevalence of drug resistance loci (Figure 6D). The emergence of C1 and C2 clades have been shaped by the acquisition of specific F-plasmids involving a series of plasmid gene gain and losses and the exchange of gene modules mediated by the insertion sequence IS26 (41). The increased rate of gain and loss in these two clades is consistent with them acquiring and exchanging multi-drug resistant (MDR) plasmids more easily than Clade A which has also undergone a recent expansion but is generally more susceptible to different classes of antibiotics (40).

**Fig. 6.**
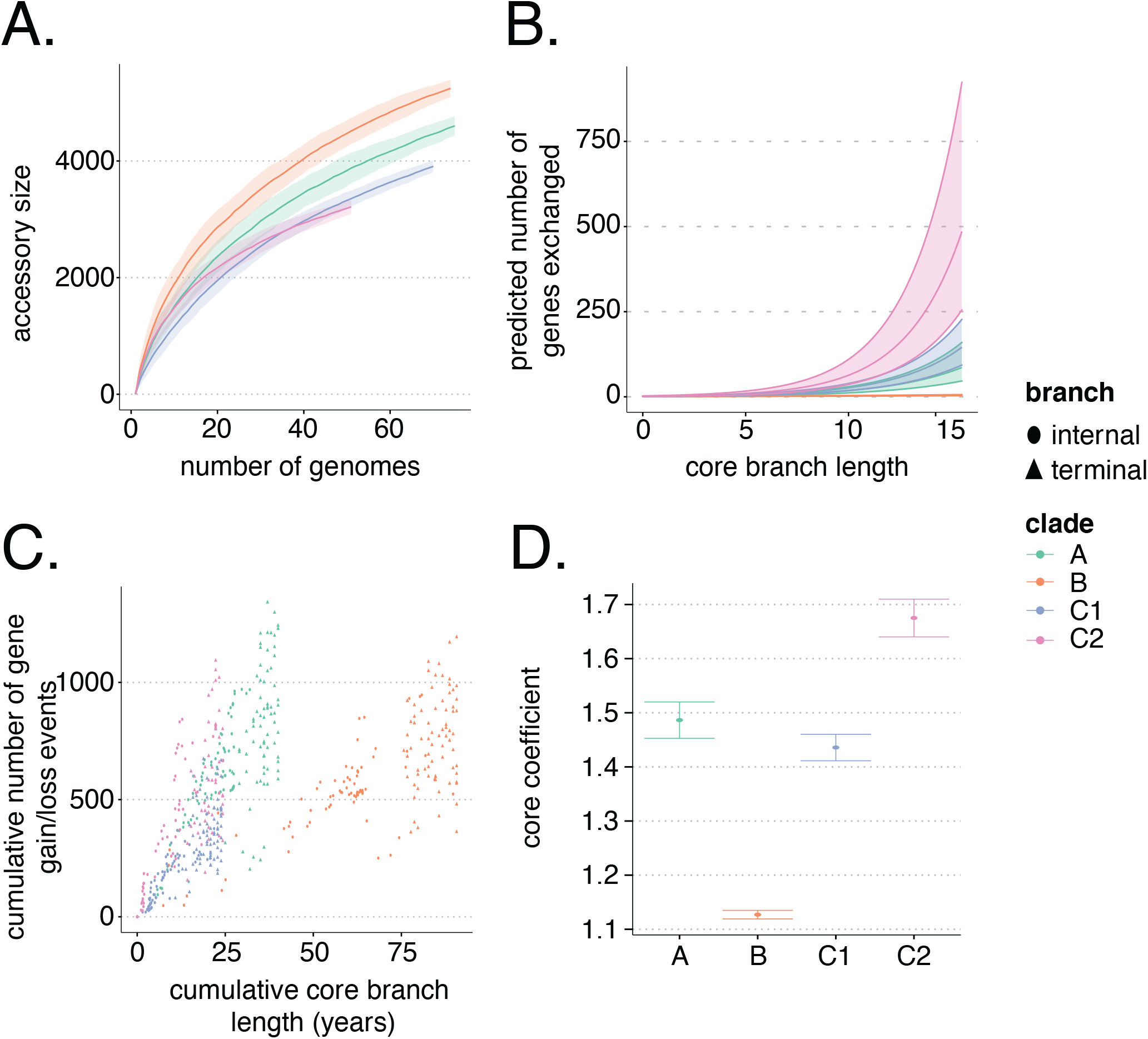
**A.)** pangenome accumulation curves for the clades in the *E. coli*-ST131 datasets. Pangenomes datasets were taken from Gladstone et al., and were constructed using Panaroo. **B.)** The corresponding predicted slope of the relationship between core genome branch length and the number of gene gain and loss events as inferred by the Panstripe algorithm. **C.)** The cumulative number of gene gain and loss events versus the cumulative branch length starting from the root node of each tree in A. This is a similar plot to the common ‘root-to-tip’ plot used in phylogenetic dating. **D.)** The estimated parameters of the Generalised Linear Model used in Panstripe. Error bars represent the 95% confidence interval of the parameter estimates. Higher values of the core coefficient indicate an increased rate of gene gain and loss.

As we have dated phylogenies for both the *E. faecalis* and *E. coli* datasets it is possible to use Panstripe to compare the rates of gene gain and loss between the two species. We found that the *E. coli* clades generally had significantly higher rates of gene gain and loss than the *E. faecalis* clades. This is in agreement with the more generalistic lifestyle of *E. faecalis*. In contrast to the specialisation of each of the *E. coli* ST131 clades mediated by the gain and loss of specific MDR plasmids, there was not a significant difference between the *E. faecalis* clades and the *E. coli* clade B (41, 42). Our ability to detect smaller differences in the rates of gene exchange is limited by the large differences in the structure of the underlying phylogenies, which leads to increased uncertainty and a reduction in the statistical power to detect differences in these pairwise comparisons.

Interestingly, although we found small differences in the error rates between the *E. coli* and *E. faecalis* datasets, there was a large difference in the dispersion parameters of the Panstripe GLM. This suggests that the number of genes involved in each gain and loss event differs significantly between the two species. The different mechanisms driving HGT in the two species and the difference in the size of the MGEs could explain this difference (43). HGT in *E. coli*, is predominantly the result of phage interactions and the exchange of large F-plasmids (size on average > 100 kbp) while the total plasmid content of *E. faecalis* is generally lower (average ∼24 kbp) (37, 41). Therefore, *E. coli* has a substantial part of the accessory genome residing on large MGEs which have shaped the evolution of the subclades whereas the events of gain and loss for *E. faecalis* may involve smaller MGEs which allocate fewer genes.

### Improved estimates of associations between phenotypes and pangenome evolutionary dynamics

The GLM framework used in Panstripe allows for other covariates and phenotypes of interest to be easily incorporated into the model and tested to identify significant associations with the rate of gene gain and loss. Common covariates of interest include whether lineages are associated with particular environments such as hospitals, drug resistance or invasive disease.

Previously, we have used the Finitely Many Genes model, as implemented in Panaroo, to estimate the rates of gene gain and loss in 51 major Global Pneumococcal Sequencing Clusters (GPCSs) (12, 23, 44, 45). The correlation between the estimated rates and the invasiveness of each lineage was estimated using Spearman’s correlation coefficient. This approach identified an association between the rate of gene gain and loss and whether the lineage had a significant odds ratio of invasive disease (44). Although Panaroo significantly reduced the number of errors, this approach was still sensitive to any remaining errors in the gene presence/absence matrix. To address this, we redid the analysis using the same gene presence/absence matrix inferred by Panaroo but with the Panstripe algorithm. Similar to our previous analysis, Panstripe identified a lower rate of gene gain and loss in GP-SCs that had a significant odds ratio of severe disease (p<1e-3, Figure 7A) (12, 44, 45). This could be a signal of genome reduction which has been linked with pathogenicity across multiple divergence scales (46).

**Fig. 7.**
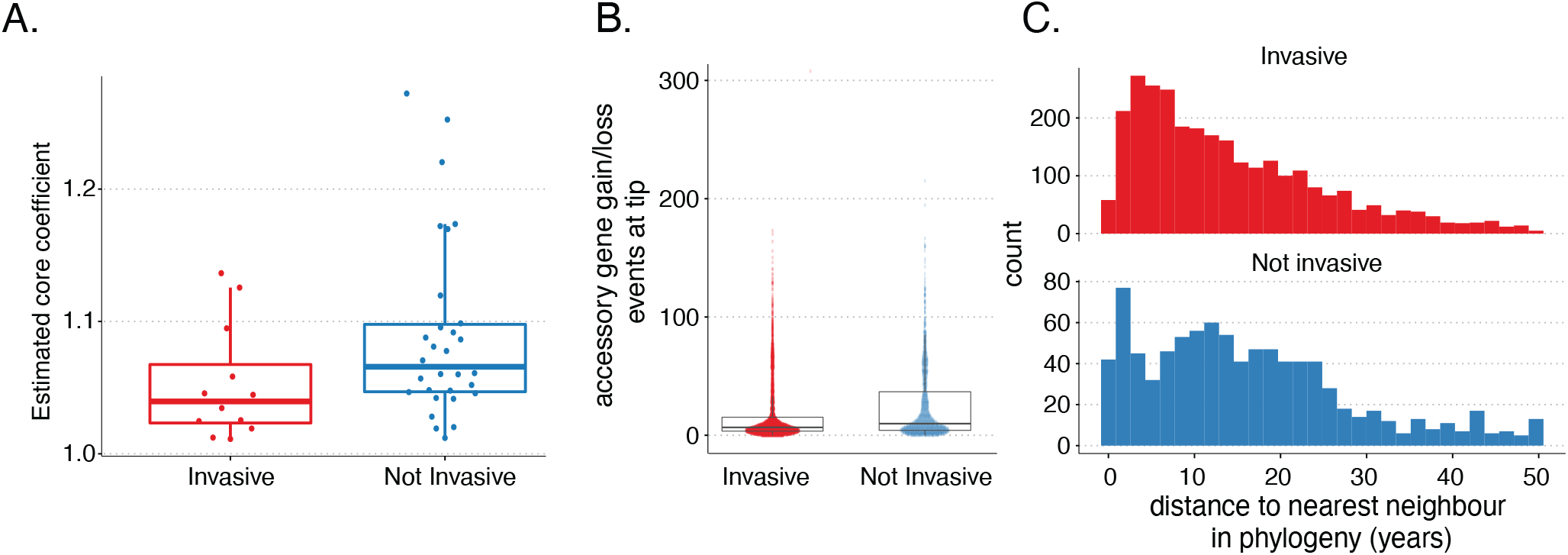
**A.)** The estimated coefficient of the core genome parameter of the Panstripe GLM for each of the GPSCs. A higher value indicates an increased rate of gene gain and loss and invasive GPSCs were found to have a decreased rate of gene exchange. **B.)** The number of accessory genes found to have been either gained or lost at the tips of the core genome phylogenies in each GPSC. **C.)** Histograms of the pairwise patristic distance (in years) between isolates in invasive and non-invasive GPSCs.

**Fig. 8.**
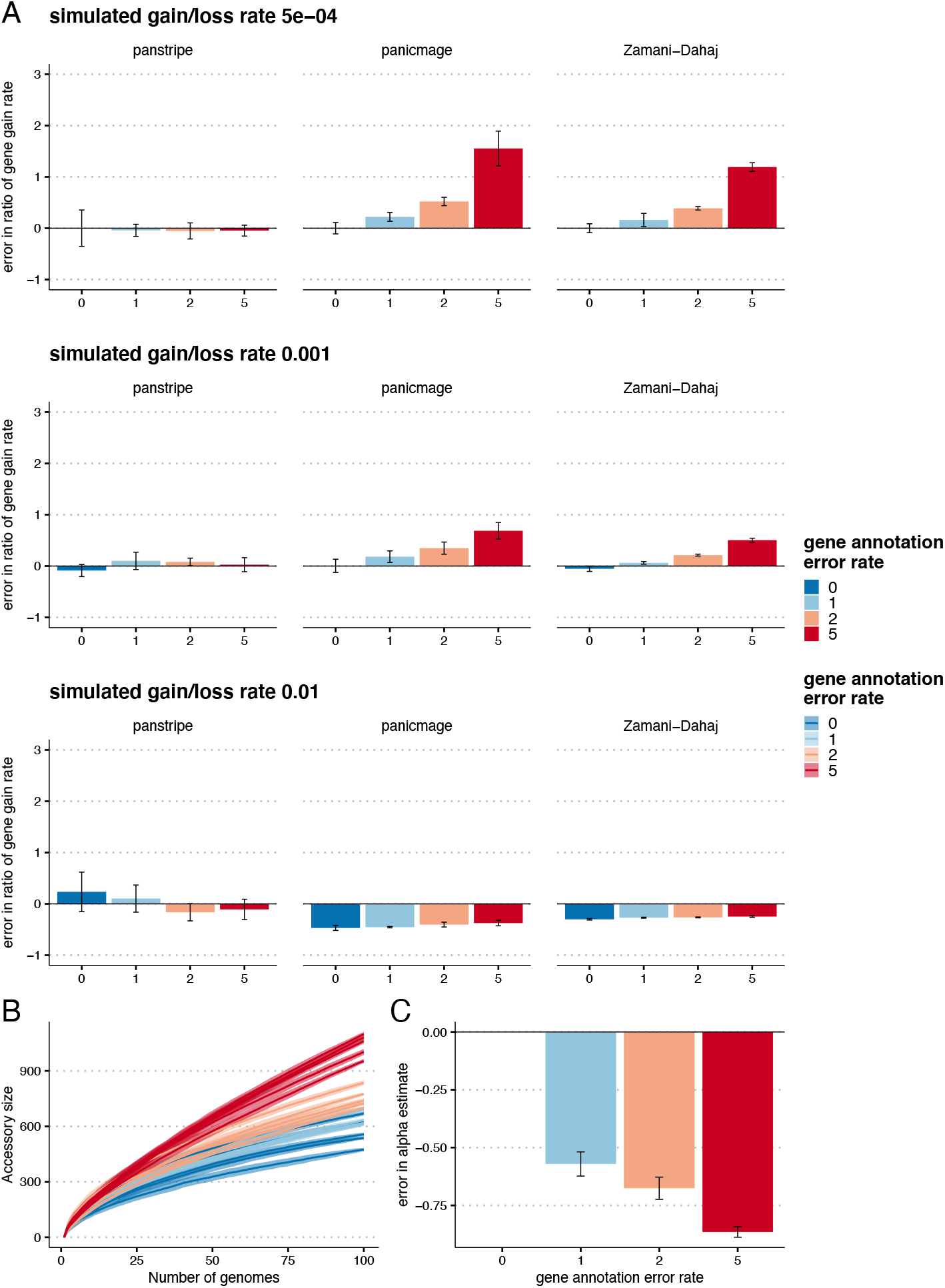
This figure reproduces the results of Figure 3 but assumes an even rate of gene gain and loss. **A.)** Bars indicate the mean percentage error in the estimated ratio of gene gain and loss rate compared to the reference rate of 5 × 10^*−*4^ with a simulated annotation error of 0. The three rows represent increasingly large gene gain and loss rates of 5 × 10^*−*4^, 0.001 and 0.01. The simulated annotation error rates are given along the x-axis. **B.)** Pangenome accumulation curves with a simulated gene gain and loss rate of 0.001. The colours represent the increasing annotation error rates. **C.)** The corresponding error in the *α* parameter estimates after fitting Heaps law to the curves in B. Lower *α* estimates indicate a more ‘open’ pangenome. Thus, higher rates of annotation error can lead to incorrect estimates of whether a pangenome is open or closed.

**Fig. 9.**
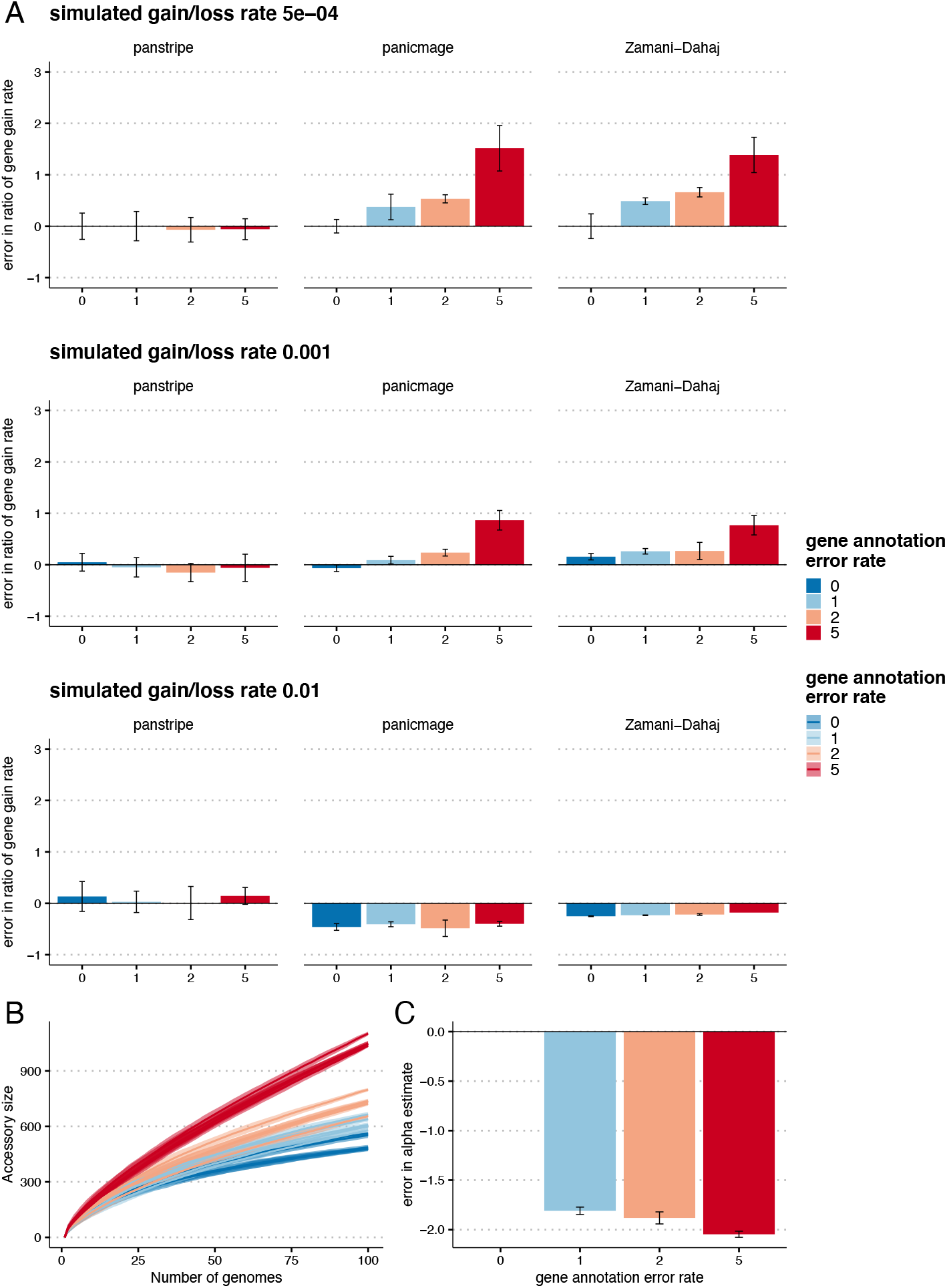
This figure reproduces the results of Figure 3 but assumes a greater rate of gene gain relative to gene loss. **A.)** Bars indicate the mean percentage error in the estimated ratio of gene gain and loss rate compared to the reference rate of 5 × 10^*−*4^ with a simulated annotation error of 0. The three rows represent increasingly large gene gain and loss rates of 5 × 10^*−*4^, 0.001 and 0.01. The simulated annotation error rates are given along the x-axis. **B.)** Pangenome accumulation curves with a simulated gene gain and loss rate of 0.001. The colours represent the increasing annotation error rates. **C.)** The corresponding error in the *α* parameter estimates after fitting Heaps law to the curves in B. Lower *α* estimates indicate a more ‘open’ pangenome. Thus, higher rates of annotation error can lead to incorrect estimates of whether a pangenome is open or closed.

Unlike our previous analysis, Panstripe found that this result was sensitive to which branches were included in the analysis with low bootstrap support for the identified association between invasiveness and gene gain and loss rate. Instead, Panstripe identified an enrichment for gene gain and loss events located on terminal branches of invasive lineages (Figure 7B). A potential explanation is that the annotation error rates differed between severe and nonsevere lineages. However, given that similar sequencing and annotation procedures were used for all genomes in the GPS project this is unlikely. Another explanation is that sampling bias or differences in the underlying population size of the lineages is driving the signal. Most invasive isolates, such as those expressing capsule serotype 1, are known to have outbreak epidemiology (44). This could lead to a greater sampling of highly related pneumococcal strains from invasive lineages. The greater evolutionary time separating non-invasive lineages would thus allow for a higher number of unique gene exchange events to occur at the tips of the phylogenies, which are more heavily influenced by faster-moving MGEs. This hypothesis is supported by looking at the pairwise patristic distance separating invasive and non-invasive isolates (Figure 7C). Here, invasive isolates are more closely related leading to a smaller number of rare unique genes being identified at the tips of the phylogeny. This highlights the utility of the Panstripe algorithm and helps to demonstrate its ability to account for annotation errors, population structure and sampling biases.

## Discussion

Determining the presence/absence of genes in prokaryotic genomes is a complex and error-prone process. Annotation and clustering errors as well as sampling bias and differences in the underlying population structure of strains and species can all complicate the analysis of pangenome dynamics. To address these problems we developed Panstripe, an algorithm which is robust to both errors in the gene presence/absence matrix and the diversity of the underlying genomes. Panstripe can compare the gene exchange rates between pangenomes and determine if these differences are associated with the underlying core genome diversity, rare and erroneous genes occurring at the tips of a phylogeny or the average size of gene gain and loss events. The use of a GLM framework also allows for associations between gene exchange and covariates of interest to be investigated.

We found that methods that do not account for the diversity of the underlying genomes perform particularly badly including the commonly used pangenome accumulation curve and Heaps power law. This was most evident in the analysis of two gene presence/absence matrices obtained after running the Panaroo and Roary pipelines on the same dataset. The pangenome accumulation curve incorrectly indicated a diverse pangenome whilst Panstripe was able to use the additional core genome phylogenetic information to correct for the large difference in error rates of the two pangenome clustering tools.

Given a lack of an association between the core genome branch length and the accessory genome it is tempting to say that such a pangenome is ‘closed’. However, the binary classification of pangenome into ‘open’ and ‘closed’ can be problematic as it does not incorporate the evolutionary timescale being considered. For example, while there is unlikely to be any gene exchange in an outbreak of *M. tuberculosis*, there is evidence of pangenome variation across the Mycobacterium genus (47–49). Instead, we suggest that it is better to report whether the core and accessory genome variation is ‘coupled’ in that there is a significant association between the core and accessory genome diversity in a particular dataset.

While we have shown that Panstripe provides considerable improvements to the analysis of pangenome dynamics, it does not implement a formal evolutionary model. Instead, we hope it provides a substantial improvement over the use of pangenome accumulation curves and allows for simple hypotheses to be tested. As more complex evolutionary models are proposed we expect that Panstripe will be used similarly to TempEst — as a check for a temporal signal before running more computationally intensive algorithms.

Panstripe is written in R and is available under the open source MIT licence from https://github.com/gtonkinhill/panstripe. The code used to generate the simulations and produce the analyses along with the gene presence/absence matrices and phylogenies is available from https://github.com/gtonkinhill/panstripe-manuscript.

## Methods

### Overview

The Panstripe algorithm takes a binary gene presence/absence matrix and an accompanying core genome phylogeny as input for each pangenome dataset being considered. Initially, the ancestral state of the presence of each gene at each node in the core genome phylogeny is inferred using either maximum parsimony or maximum likelihood methods. Using these estimates, the total number of gene gain and loss events on each branch of the phylogeny is calculated. A GLM framework is then used to compare the number of gene gain and loss events with the branch length of the core genome phylogeny. Terms are also included in the GLM to indicate the depth of the branch within the phylogeny and whether or not the branch occurs at the tips of the tree. These help to control for the reduced ability to observe gene gain and loss events at higher branches of the tree as well as errors in the gene presence/absence matrix respectively. Comparisons between pangenomes are made by including an additional categorical covariate in the GLM for the pangenome and investigating the interaction terms between this variable and the core, tip and depth terms.

### Ancestral state reconstruction

Panstripe includes two methods for inferring whether a gene was present or not at each ancestral node of the core genome phylogeny. The default maximum parsimony approach uses a version of Sankoff’s dynamic programming algorithm adapted from the Castor R package to determine the ancestral states that correspond with the smallest number of state changes along the phylogeny (27, 28). Alternatively, either a maximum likelihood based method or stochastic mapping can be used. The included maximum-likelihood based method assumes a fixed-rates continuous-time Markov model (Mk model) as implemented in the Ape R package (29, 50). The simulation-free version of stochastic mapping is included as implemented in the Sfreemap R package **??**. After reconstructing the ancestral state for each gene independently the total number of gene gain and loss events for each branch is found by taking the sum of events for each gene assuming that at most one change occurs between consecutive nodes.

While maximum likelihood based methods can provide improved estimates of the ancestral states in some instances (51), using a maximum-likelihood method effectively uses the branch lengths of the core genome phylogeny twice. Once in the ancestral state reconstruction and once in the regression. This can artificially inflate the correlation between the branch lengths and the number of gene gain and loss events. Consequently, we prefer to use maximum-parsimony. In practice, we have found that using either approach gives very similar results in most instances.

### Tweedie generalised linear regression model

After inferring the number of gene gain and loss events, each branch of the phylogeny is subsequently treated as an independent data point. For each branch *i*, let *g*_*i*_ be the number of gene gain and loss events, *l*_*i*_ the core branch length, *d*_*i*_ the core genetic distance from the root of the phylogeny to the node at the start of the branch, and *k*_*i*_ be a binary variable indicating whether or not the branch occurred at the tip of the phylogeny.

We assume that the number of gene gain and loss events *g*_*i*_ follows a compound Poisson distribution. That is, the number of HGT events follows a Poisson distribution but the size of each event follows a Gamma distribution. This is important as HGT events frequently involve the acquisition or loss of multiple genes. To estimate the association between the number of genes gained and lost and the other covariates we make use of a Tweedie Generalised Linear Model implemented in the Statmod R package (52). The Tweedie distribution generalises many distributions commonly used in GLMs including the Normal, Poisson and Gamma distributions depending upon the index power parameter *p*. For 1 *< p <* 2, the Tweedie distribution is equivalent to a compound Poisson distribution (53).

A Tweedie GLM with a log link function assumes that the expectation of the *i*th response *µ*_*i*_ := *E*(*g*_*i*_) is related to a vector *x*_*i*_ of covariates with corresponding coefficients *b* by

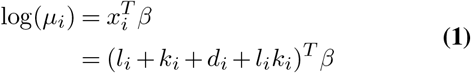

The corresponding variance of *g*_*i*_ is given by

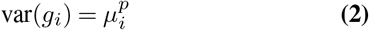

We include an interaction term between the branch length and whether a branch is terminal to account for the increased chance that a gene is both gained and erroneously omitted on longer terminal branches. For datasets with reliable phylogenies, it is generally safe to assume that annotation errors will be unlikely to propagate to non-terminal branches during ancestral state reconstruction. The ability of the algorithm to accurately characterise the gene gain and loss rate can then be improved by setting the intercept of the GLM to zero. This effectively assumes that no gene gain and loss events can occur on internal branches of zero length. If a phylogeny is thought to be unreliable an intercept can be included to account for the mean error of inferred gain and loss events on internal branches being non-zero. This has the side effect of reducing the sensitivity of the approach to detect differences between pangenome datasets. Panstripe allows the user to choose which approach is best suited for their analysis with the intercept excluded by default.

Using this framework, it is possible to perform standard hypothesis tests based on a Student t-distribution to determine whether each of the parameters is significantly associated with gene gain and loss. A significant association with the core branch length indicates that there is evidence of HGT at the upper branches of the phylogeny. The association with the ‘tip’ covariate can be driven by a combination of both the error rate in the inference of the gene presence/absence matrix and the acquisition and loss of rare genes that are seen only once. The depth parameter accounts for the potential loss in power to identify HGT at higher branches of the phylogeny.

### Comparing pangenomes

Panstripe uses interaction terms to compare the relationship of the covariates with gene gain and loss between two pangenome datasets. While the use of a standard GLM framework would allow for the comparison of the inferred slope or *β* parameters between datasets, it assumes a single dispersion parameter. This effectively fixes the relationship between the rate of gene exchange and the size of each HGT event. This can be seen in the relationship between the Tweedie distribution parameters and the corresponding parameters of the Poisson and Gamma components of the Compound Poisson distribution

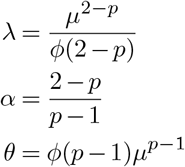

Here, *λ* is the mean of the Poisson distribution and *α* and *θ* are the shape and scale parameters of the Gamma distribution respectively.

To relax this assumption, Panstripe optionally allows for the dispersion parameter to vary between pangenomes using the Double Generalised Linear Model (DGLM) framework described in Smyth, (32). The DGLM framework models the mean and dispersion using two separate GLMs with both being a function of the covariates. A maximum likelihood estimate of the parameters is then found by alternating between the two sub-models. Setting the pangenome being considered for branch *i* as a binary covariate *n*_*i*_ the DGLM used to compare pangenomes can be formulated as

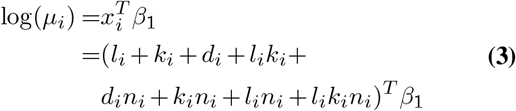

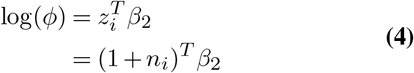

The significance of the inferred coefficients of the interaction terms can be used to determine whether the association between HGT and the depth, core and tip covariates differs significantly between pangenomes. When investigating the dispersion parameter, Panstripe uses the Likelihood Ratio Test to determine whether there is a significant benefit to accounting for a variable dispersion between two pangenome datasets. In some cases, such as when comparing lineages from the same species, it may be safe to assume that the relationship between the mean and dispersion is similar in both pangenomes. In this case, it can be preferable to fix the dispersion parameter to improve the statistical power of an analysis.

### Identifying associations with additional covariates

The GLM framework used by Panstripe allows for other covariates that may influence pangenome dynamics to be considered. This could include whether or not a lineage is highly drug-resistant, hospital-associated or frequently linked to invasive disease. In general, this is most useful when many separate lineages from a large genome collection of a single species are available as is becoming increasingly common with the advent of large sequencing studies and the introduction of WGS as a routine service for public health surveillance (44, 45, 54). To investigate such associations, Panstripe replaces the binary pangenome covariate in equation 3 with the covariate(s) of interest. These can be logical, continuous or categorical.

### Bootstrap confidence intervals

Panstripe estimates confidence intervals for each coefficient using the Bootstrap, by resampling each branch with replacement (55). While it is possible to use the GLM model to obtain confidence interval estimates, this makes some assumptions about the distribution of the coefficients and is more susceptible to outliers. Panstripe uses the boot R package to calculate these intervals which implements several commonly used methods for calculating the interval from the resampled set of coefficients (55, 56). In general, we have found the Bootstrap confidence intervals to provide similar estimates to that of the GLM model. However, in some cases, differences have indicated a high sensitivity of the result to particular branches. Optionally, Panstripe also includes the possibility of estimating Bootstrap p-values using the confidence interval inversion method (57).

### Simulations

To compare the performance of Panstripe we simulated core genome phylogenies using the ‘rtree’ function from the ape package (58). The number of gene gain and loss events on each branch was simulated using the simSeq function in the Phangorn package (59) with an elevated frequency of gene loss events (base frequency vector of [0.3, 0.2]). To verify that Panstripe is robust to different gene gain/loss ratios we repeated the analysis with both equal base frequencies and an elevated rate of gene gain events. We assume that each gain or loss event will involve the same set of genes and simulate the size of the set using a Poisson distribution as implemented in R. Errors in the pangenome matrix were simulated by randomly adding or removing single entries in the final gene presence/absence matrix. The number of errors for each genome was simulated using a Poisson distribution. The full set of parameters used in the simulations is given in Supplementary Table **??**. All code to reproduce the analyses is available in the accompanying GitHub repository.

### Dataset preparation

The phylogenies and pangenome gene presence-absence matrices for the *E. coli* and *E. faecalis* datasets were taken directly from the respective publications (37, 40). Briefly, in both datasets clades were defined via alignment-free whole-genome clustering using Population Partitioning Using Nucleotide K-mers (PopPUNK) v.1.2.2 (60). Phylogenies of the resulting clades were then constructed using RaxML v8.2.8 with GTR + Gamma rate model after removing recombination using Gubbins v2.4.0. Pangenomes for both datasets were generated using Panaroo v1.2. A more detailed description of the methods can be found in the original publications.

The pangenome of the *M. tuberculosis* dataset was taken directly from the publication of the Panaroo algorithm (Panaroo v1.0.0). This included 414 genomes which were very closely related, originally published as part of an analysis into an outbreak of isoniazid-resistant tuberculosis in London (33). As our pangenome dataset included some genomes that were not included in the phylogeny of the original Mtb publication, we reconstructed a new phylogeny. This was achieved by generating a core genome from the original assemblies using Snippy v4.6. Gubbins v2.4.0 was then used to remove poor-quality regions of the alignment and the final phylogeny was generated using Fasttree v2.1.11.

## Data and materials availability

Panstripe is available under an MIT open source licence on GitHub https://github.com/gtonkinhill/panstripe (archived at 10.5281/zenodo.6404363). The code, phylogenies and gene presence/absence matrices used in our analyses are available at https://github.com/gtonkinhill/panstripe-manuscript (archived at 10.5281/zenodo.6404403).

## Acknowledgements

We thank the members of the Davies group at the Doherty Institute for helpful discussions and testing of the software.

## Supplementary Figures

**Supplementary Figure 1.**
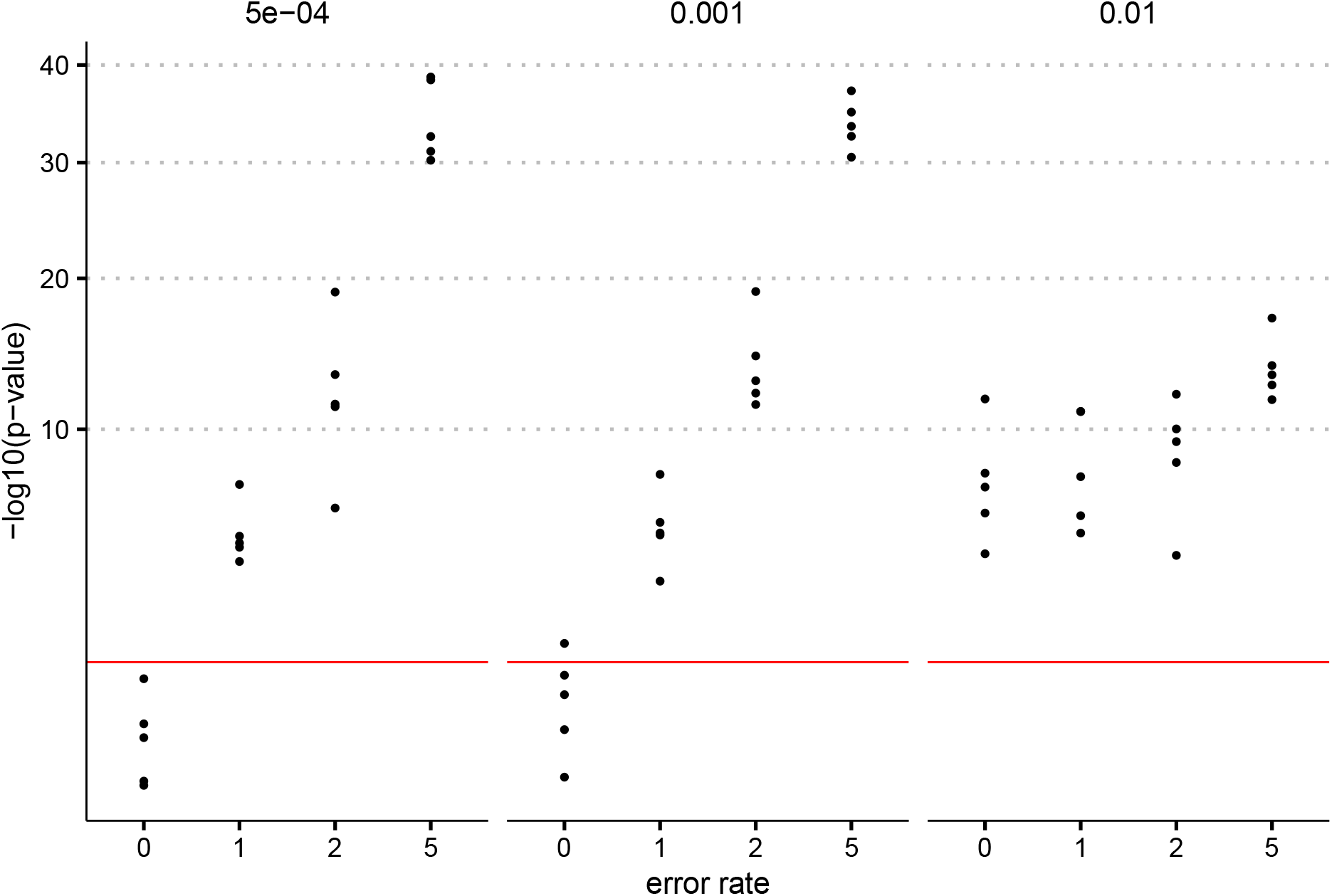
P-values for the covariate indicating whether or not a branch occurred at the tip of a phylogeny. Five different error rates were simulated along with four symmetric gene gain/loss rates. Five replicates were simulated for each parameter set. The top row of plots indicates the resulting p-values when comparing each dataset with a pangenome with a mean error rate of 1 while the bottom row shows comparisons with an error rate of 5. The horizontal red line indicates a p-value of 0.05 and points above this line are considered significantly different. Panstripe was less sensitive to differences in error rates when the gene gain and loss rate was very high (*≥* 1*e −* 2) which is rarely seen in practice.

**Supplementary Figure 2.**
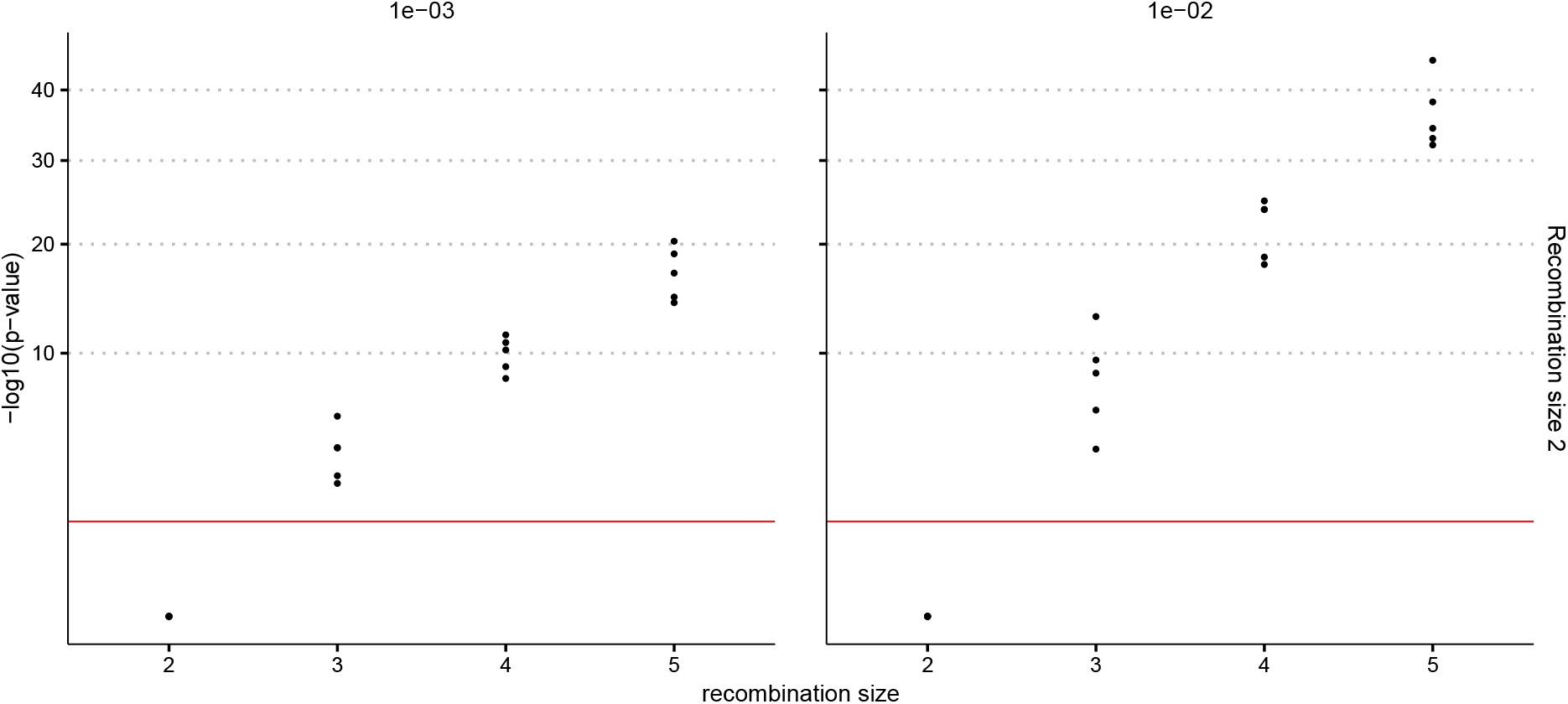
P-values obtained after performing a Likelihood Ratio Test to compare a model which allowed the dispersion parameter to vary between pangenomes with a model with a fixed dispersion parameter. The mean number of genes involved in a recombination event was simulated for five different parameter values and each was compared with a mean size of 2. The horizontal red line indicates a p-value of 0.05 and points above this line are considered significantly different.

**Supplementary Figure 3.**
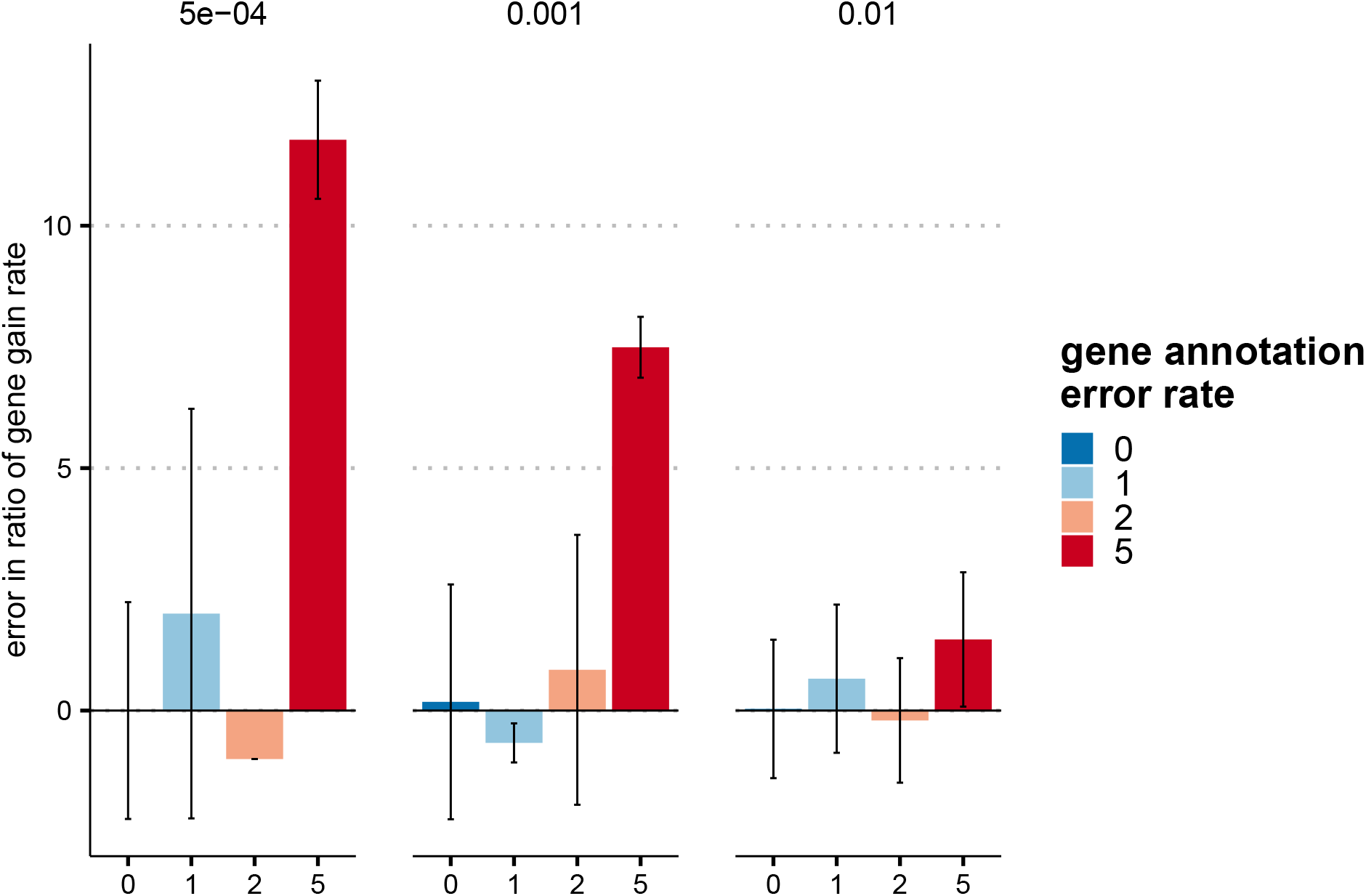
Bars indicate the mean percentage error in the estimated ratio of gene gain and loss rate compared to the reference rate of 5 × 10^*−*4^ with a simulated annotation error of 0. The R code of Collins et al. gave very inconsistent results as indicated by the very large standard deviation in the estimated ratio (error bars) that often span multiple orders of magnitude. This inconsistency makes it challenging to compare results between runs using this method.

**Supplementary Figure 4.**
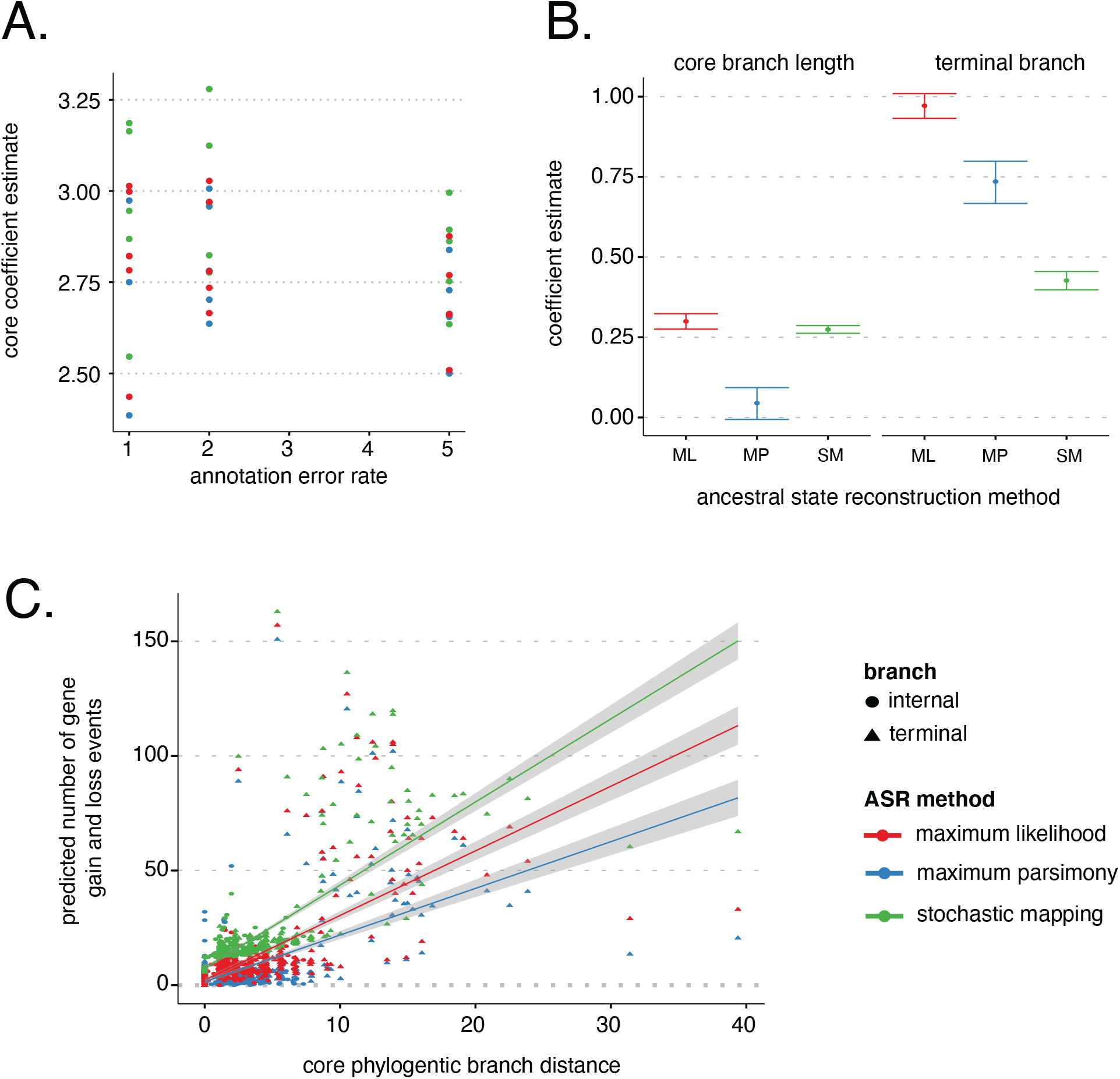
**A.)** The estimated core coefficient of the Panstripe GLM after running the algorithm on simulated pangenomes using a gene gain and loss rate of 1e-3 and three different gene annotation error rates (1, 2 and 5). The colours of the points indicate the ancestral state reconstruction algorithm used. The higher variation within each error rate indicates that the Panstripe algorithm is robust to the choice of ASR algorithm on relatively well-behaved datasets. **B.)** The estimated core and tip coefficients of the Panstripe GLM after running the algorithm on the highly clonal Mtb dataset. Here, the very low core genome diversity made constructing an accurate phylogeny difficult. The resulting phylogeny had very short branch lengths and several multichotomies that led to greater discrepancies between the results of each ASR algorithm. Maximum parsimony, which ignores branch lengths, provided the most reliable result indicating a core coefficient that was not significantly different from zero. **C.)** A dot plot indicating the inferred core genome branch lengths versus the corresponding number of gene gain and loss events inferred to have occurred on each branch using the three ASR algorithms. The lines represent the best fit of a linear model and help to highlight the average differences between the three ASR methods.

**Supplementary Table 1.**
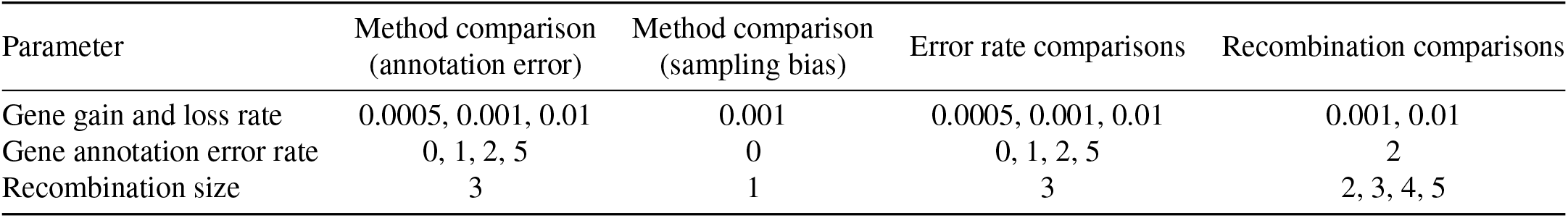
The parameters used to compare the results of each method on the simulated pangenome datasets.

## References

1. Christopher M Thomas and Kaare M Nielsen. Mechanisms of, and barriers to, horizontal gene transfer between bacteria. Nat. Rev. Microbiol., 3(9):711–721, September 2005.

2. Brian J Arnold, I-Ting Huang, and William P Hanage. Horizontal gene transfer and adaptive evolution in bacteria. Nat. Rev. Microbiol., November 2021.

3. D Dubnau. DNA uptake in bacteria. Annu. Rev. Microbiol, 53:217–244, 1999.

4. Simon R Harris, Edward J Feil, Matthew T G Holden, Michael A Quail, Emma K Nickerson, Narisara Chantratita, Susana Gardete, Ana Tavares, Nick Day, Jodi A Lindsay, Jonathan D Edgeworth, Hermínia de Lencastre, Julian Parkhill, Sharon J Peacock, and Stephen D Bentley. Evolution of MRSA during hospital transmission and intercontinental spread. Science, 327(5964):469–474, January 2010.

5. Nicholas J Croucher, Jonathan A Finkelstein, Stephen I Pelton, Patrick K Mitchell, Grace M Lee, Julian Parkhill, Stephen D Bentley, William P Hanage, and Marc Lipsitch. Population genomics of post-vaccine changes in pneumococcal epidemiology. Nat. Genet., 45(6):656–663, June 2013.

6. Kelly L Wyres, Ryan R Wick, Louise M Judd, Roni Froumine, Alex Tokolyi, Claire L Gorrie, Margaret M C Lam, Sebastián Duchêne, Adam Jenney, and Kathryn E Holt. Distinct evolutionary dynamics of horizontal gene transfer in drug resistant and virulent clones of klebsiella pneumoniae. PLoS Genet., 15(4):e1008114, April 2019.

7. Duccio Medini, Claudio Donati, Hervé Tettelin, Vega Masignani, and Rino Rappuoli. The microbial pan-genome. Curr. Opin. Genet. Dev., 15(6):589–594, December 2005.

8. Wei Ding, Franz Baumdicker, and Richard A Neher. panx: pan-genome analysis and exploration. Nucleic Acids Res., 46(1):e5, January 2018.

9. Guillaume Gautreau, Adelme Bazin, Mathieu Gachet, Rémi Planel, Laura Burlot, Mathieu Dubois, Amandine Perrin, Claudine Médigue, Alexandra Calteau, Stéphane Cruveiller, Catherine Matias, Christophe Ambroise, Eduardo P C Rocha, and David Vallenet. PPanG-GOLiN: Depicting microbial diversity via a partitioned pangenome graph. PLoS Comput. Biol., 16(3):e1007732, March 2020.

10. Andrew J Page, Carla A Cummins, Martin Hunt, Vanessa K Wong, Sandra Reuter, Matthew T G Holden, Maria Fookes, Daniel Falush, Jacqueline A Keane, and Julian Parkhill. Roary: rapid large-scale prokaryote pan genome analysis. Bioinformatics, 31(22):3691–3693, November 2015.

11. Sion C Bayliss, Harry A Thorpe, Nicola M Coyle, Samuel K Sheppard, and Edward J Feil. PIRATE: A fast and scalable pangenomics toolbox for clustering diverged orthologues in bacteria. Gigascience, 8(10), October 2019.

12. Gerry Tonkin-Hill, Neil MacAlasdair, Christopher Ruis, Aaron Weimann, Gal Horesh, John A Lees, Rebecca A Gladstone, Stephanie Lo, Christopher Beaudoin, R Andres Floto, Simon D W Frost, Jukka Corander, Stephen D Bentley, and Julian Parkhill. Producing polished prokaryotic pangenomes with the panaroo pipeline. Genome Biol., 21(1):180, July 2020.

13. Zhemin Zhou, Jane Charlesworth, and Mark Achtman. Accurate reconstruction of bacterial pan-and core genomes with PEPPAN. Genome Res., 30(11):1667–1679, November 2020.

14. Karl I Ugland, John S Gray, and Kari E Ellingsen. The species-accumulation curve and estimation of species richness. J. Anim. Ecol., 72(5):888–897, September 2003.

15. Hervé Tettelin, David Riley, Ciro Cattuto, and Duccio Medini. Comparative genomics: the bacterial pan-genome. Curr. Opin. Microbiol., 11(5):472–477, October 2008.

16. Boris G Mirkin, Trevor I Fenner, Michael Y Galperin, and Eugene V Koonin. Algorithms for computing parsimonious evolutionary scenarios for genome evolution, the last universal common ancestor and dominance of horizontal gene transfer in the evolution of prokaryotes. BMC Evol. Biol., 3:2, January 2003.

17. Ofir Cohen and Tal Pupko. Inference and characterization of horizontally transferred gene families using stochastic mapping. Mol. Biol. Evol., 27(3):703–713, March 2010.

18. Weilong Hao and G Brian Golding. The fate of laterally transferred genes: life in the fast lane to adaptation or death. Genome Res., 16(5):636–643, May 2006.

19. Mira V Han, Gregg W C Thomas, Jose Lugo-Martinez, and Matthew W Hahn. Estimating gene gain and loss rates in the presence of error in genome assembly and annotation using CAFE 3. Mol. Biol. Evol., 30(8):1987–1997, August 2013.

20. Liang Liu, Lili Yu, Venugopal Kalavacharla, and Zhanji Liu. A bayesian model for gene family evolution. BMC Bioinformatics, 12:426, November 2011.

21. F Baumdicker, W R Hess, and P Pfaffelhuber. The diversity of a distributed genome in bacterial populations. Ann. Appl. Probab., 20(5):1567–1606, 2010. ISSN 1050-5164.

22. Franz Baumdicker, Wolfgang R Hess, and Peter Pfaffelhuber. The infinitely many genes model for the distributed genome of bacteria. Genome Biol. Evol., 4(4):443–456, February 2012.

23. Seyed Alireza Zamani-Dahaj, Mohamed Okasha, Jakub Kosakowski, and Paul G Higgs. Estimating the frequency of horizontal gene transfer using phylogenetic models of gene gain and loss. Mol. Biol. Evol., 33(7):1843–1857, July 2016.

24. R Eric Collins and Paul G Higgs. Testing the infinitely many genes model for the evolution of the bacterial core genome and pangenome. Mol. Biol. Evol., 29(11):3413–3425, November 2012.

25. Steven L Salzberg. Next-generation genome annotation: we still struggle to get it right. Genome Biol., 20(1):92, May 2019.

26. John A Lees, Michelle Kendall, Julian Parkhill, Caroline Colijn, Stephen D Bentley, and Simon R Harris. Evaluation of phylogenetic reconstruction methods using bacterial whole genomes: a simulation based study. Wellcome Open Res, 3:33, March 2018.

27. David Sankoff. Minimal mutation trees of sequences. SIAM J. Appl. Math., 28(1):35–42, January 1975.

28. Stilianos Louca and Michael Doebeli. Efficient comparative phylogenetics on large trees. Bioinformatics, 34(6):1053–1055, March 2018.

29. Z Yang, S Kumar, and M Nei. A new method of inference of ancestral nucleotide and amino acid sequences. Genetics, 141(4):1641–1650, December 1995.

30. Andrew Rambaut, Tommy T Lam, Luiz Max Carvalho, and Oliver G Pybus. Exploring the temporal structure of heterochronous sequences using TempEst (formerly Path-O-Gen). Virus Evol, 2(1):vew007, January 2016.

31. Alexei J Drummond, Oliver G Pybus, Andrew Rambaut, Roald Forsberg, and Allen G Rodrigo. Measurably evolving populations. Trends Ecol. Evol., 18(9):481–488, September 2003.

32. Gordon K Smyth. Generalized linear models with varying dispersion. J. R. Stat. Soc., 51 (1):47–60, September 1989.

33. Nicola Casali, Agnieszka Broda, Simon R Harris, Julian Parkhill, Timothy Brown, and Francis Drobniewski. Whole genome sequence analysis of a large Isoniazid-Resistant tuberculosis outbreak in london: A retrospective observational study. PLoS Med., 13(10):e1002137, October 2016.

34. Hervé Tettelin, Vega Masignani, Michael J Cieslewicz, Claudio Donati, Duccio Medini, Naomi L Ward, Samuel V Angiuoli, Jonathan Crabtree, Amanda L Jones, A Scott Durkin, Robert T Deboy, Tanja M Davidsen, Marirosa Mora, Maria Scarselli, Immaculada Margarit y Ros, Jeremy D Peterson, Christopher R Hauser, Jaideep P Sundaram, William C Nelson, Ramana Madupu, Lauren M Brinkac, Robert J Dodson, Mary J Rosovitz, Steven A Sullivan, Sean C Daugherty, Daniel H Haft, Jeremy Selengut, Michelle L Gwinn, Liwei Zhou, Nikhat Zafar, Hoda Khouri, Diana Radune, George Dimitrov, Kisha Watkins, Kevin J B O’Connor, Shannon Smith, Teresa R Utterback, Owen White, Craig E Rubens, Guido Grandi, Lawrence C Madoff, Dennis L Kasper, John L Telford, Michael R Wessels, Rino Rappuoli, and Claire M Fraser. Genome analysis of multiple pathogenic isolates of streptococcus agalactiae: implications for the microbial “pan-genome”. Proc. Natl. Acad. Sci. U. S. A., 102(39):13950–13955, September 2005.

35. James S Farris. Phylogenetic analysis under dollo’s law. Syst. Zool., 26(1):77–88, 1977. ISSN 0039-7989. doi: 10.2307/2412867.

36. Katinka J Apagyi, Christophe Fraser, and Nicholas J Croucher. Transformation asymmetry and the evolution of the bacterial accessory genome. Mol. Biol. Evol., 35(3):575–581, March 2018. ISSN 0737-4038, 1537-1719. doi: 10.1093/molbev/msx309.

37. Anna K Pöntinen, Janetta Top, Sergio Arredondo-Alonso, Gerry Tonkin-Hill, Ana R Freitas, Carla Novais, Rebecca A Gladstone, Maiju Pesonen, Rodrigo Meneses, Henri Pesonen, John A Lees, Dorota Jamrozy, Stephen D Bentley, Val F Lanza, Carmen Torres, Luisa Peixe, Teresa M Coque, Julian Parkhill, Anita C Schürch, Rob J L Willems, and Jukka Corander. Apparent nosocomial adaptation of enterococcus faecalis predates the modern hospital era. Nat. Commun., 12(1):1–13, March 2021.

38. Kelli L Palmer, Paul Godfrey, Allison Griggs, Veronica N Kos, Jeremy Zucker, Christopher Desjardins, Gustavo Cerqueira, Dirk Gevers, Suzanne Walker, Jennifer Wortman, Michael Feldgarden, Brian Haas, Bruce Birren, and Michael S Gilmore. Comparative genomics of enterococci: variation in enterococcus faecalis, clade structure in e. faecium, and defining characteristics of e. gallinarum and e. casseliflavus. MBio, 3(1):e00318–11, March 2012.

39. Bernd Neumann, Karola Prior, Jennifer K Bender, Dag Harmsen, Ingo Klare, Stephan Fuchs, Astrid Bethe, Daniela Zühlke, André Göhler, Stefan Schwarz, Kirsten Schaffer, Katharina Riedel, Lothar H Wieler, and Guido Werner. A core genome multilocus sequence typing scheme for enterococcus faecalis. J. Clin. Microbiol., 57(3), March 2019.

40. Rebecca A Gladstone, Alan McNally, Anna K Pöntinen, Gerry Tonkin-Hill, John A Lees, Kusti Skytén, François Cléon, Martin O K Christensen, Bjørg C Haldorsen, Kristina K Bye, Karianne W Gammelsrud, Reidar Hjetland, Angela Kümmel, Hege E Larsen, Paul Christoffer Lindemann, Iren H Löhr, Åshild Marvik, Einar Nilsen, Marie T Noer, Gunnar S Simonsen, Martin Steinbakk, Ståle Tofteland, Marit Vattøy, Stephen D Bentley, Nicholas J Croucher, Julian Parkhill, Pål J Johnsen, Ørjan Samuelsen, and Jukka Corander. Emergence and dissemination of antimicrobial resistance in escherichia coli causing bloodstream infections in norway in 2002–17: a nationwide, longitudinal, microbial population genomic study. The Lancet Microbe, May 2021.

41. Timothy J Johnson, Jessica L Danzeisen, Bonnie Youmans, Kyle Case, Katharine Llop, Jeannette Munoz-Aguayo, Cristian Flores-Figueroa, Maliha Aziz, Nicole Stoesser, Evgeni Sokurenko, Lance B Price, and James R Johnson. Separate F-Type plasmids have shaped the evolution of the H30 subclone of escherichia coli sequence type 131. mSphere, 1(4), July 2016.

42. Kira Kondratyeva, Mali Salmon-Divon, and Shiri Navon-Venezia. Meta-analysis of pandemic escherichia coli ST131 plasmidome proves restricted plasmid-clade associations. Sci. Rep., 10(1):36, January 2020.

43. Michael A Brockhurst, Ellie Harrison, James P J Hall, Thomas Richards, Alan McNally, and Craig MacLean. The ecology and evolution of pangenomes. Curr. Biol., 29(20):R1094–R1103, October 2019.

44. Rebecca A Gladstone, Stephanie W Lo, John A Lees, Nicholas J Croucher, Andries J van Tonder, Jukka Corander, Andrew J Page, Pekka Marttinen, Leon J Bentley, Theresa J Ochoa, Pak Leung Ho, Mignon du Plessis, Jennifer E Cornick, Brenda Kwambana-Adams, Rachel Benisty, Susan A Nzenze, Shabir A Madhi, Paulina A Hawkins, Dean B Everett, Martin Antonio, Ron Dagan, Keith P Klugman, Anne von Gottberg, Lesley McGee, Robert F Breiman, and Stephen D Bentley. International genomic definition of pneumococcal lineages, to contextualise disease, antibiotic resistance and vaccine impact. EBioMedicine, 43:338–346, May 2019.

45. Rebecca A Gladstone, Stephanie W Lo, Richard Goater, Corin Yeats, Ben Taylor, James Hadfield, John A Lees, Nicholas J Croucher, Andries J van Tonder, Leon J Bentley, Fu Xi-ang Quah, Anne J Blaschke, Nicole L Pershing, Carrie L Byington, Veeraraghavan Balaji, Waleria Hryniewicz, Betuel Sigauque, K L Ravikumar, Samanta Cristine Grassi Almeida, Theresa J Ochoa, Pak Leung Ho, Mignon du Plessis, Kedibone M Ndlangisa, Jennifer E Cornick, Brenda Kwambana-Adams, Rachel Benisty, Susan A Nzenze, Shabir A Madhi, Paulina A Hawkins, Andrew J Pollard, Dean B Everett, Martin Antonio, Ron Dagan, Keith P Klugman, Anne von Gottberg, Benjamin J Metcalf, Yuan Li, Bernard W Beall, Lesley McGee, Robert F Breiman, David M Aanensen, Stephen D Bentley, and The Global Pneumococcal Sequencing Consortium. Visualizing variation within global pneumococcal sequence clusters (GPSCs) and country population snapshots to contextualize pneumococcal isolates. Microbial Genomics, 6(5), May 2020.

46. Gemma G R Murray, Jane Charlesworth, Eric L Miller, Michael J Casey, Catrin T Lloyd, Marcelo Gottschalk, Alexander W Dan Tucker, John J Welch, and Lucy A Weinert. Genome reduction is associated with bacterial pathogenicity across different scales of temporal and ecological divergence. Mol. Biol. Evol., 38(4):1570–1579, April 2021.

47. Jennifer Becq, Maria Cristina Gutierrez, Vania Rosas-Magallanes, Jean Rauzier, Brigitte Gicquel, Olivier Neyrolles, and Patrick Deschavanne. Contribution of horizontally acquired genomic islands to the evolution of the tubercle bacilli. Mol. Biol. Evol., 24(8):1861–1871, August 2007.

48. Vania Rosas-Magallanes, Patrick Deschavanne, Lluis Quintana-Murci, Roland Brosch, Brigitte Gicquel, and Olivier Neyrolles. Horizontal transfer of a virulence operon to the ancestor of mycobacterium tuberculosis. Mol. Biol. Evol., 23(6):1129–1135, June 2006.

49. Á Chiner-Oms, L Sánchez-Busó, J Corander, S Gagneux, S R Harris, D Young, F González-Candelas, and I Comas. Genomic determinants of speciation and spread of the mycobacterium tuberculosis complex. Sci Adv, 5(6):eaaw3307, June 2019.

50. Emmanuel Paradis, Julien Claude, and Korbinian Strimmer. APE: Analyses of phylogenetics and evolution in R language. Bioinformatics, 20(2):289–290, January 2004. ISSN 1367-4803. doi: 10.1093/bioinformatics/btg412.

51. Mark Pagel. The maximum likelihood approach to reconstructing ancestral character states of discrete characters on phylogenies. Syst. Biol., 48(3):612–622, 1999.

52. Peter K Dunn and Gordon K Smyth. Generalized Linear Models With Examples in R. Springer, New York, NY, 2018.

53. Bent Jørgensen. Exponential dispersion models. J. R. Stat. Soc., 49(2):127–145, January 1987.

54. Marie Anne Chattaway, Timothy J Dallman, Lesley Larkin, Satheesh Nair, Jacquelyn Mc-Cormick, Amy Mikhail, Hassan Hartman, Gauri Godbole, David Powell, Martin Day, Robert Smith, and Kathie Grant. The transformation of reference microbiology methods and surveillance for salmonella with the use of whole genome sequencing in england and wales. Front Public Health, 7:317, November 2019.

55. Bradley Efron. Better bootstrap confidence intervals. J. Am. Stat. Assoc., 82(397):171–185, March 1987.

56. A C Davison and D V Hinkley. Bootstrap Methods and Their Application. Cambridge University Press, October 1997.

57. Peter Hall. The Bootstrap and Edgeworth Expansion. Springer Science & Business Media, December 2013.

58. Emmanuel Paradis. Analysis of Phylogenetics and Evolution with R. Springer, New York, NY, 2006.

59. Klaus Peter Schliep. phangorn: phylogenetic analysis in R. Bioinformatics, 27(4):592–593, February 2011.

60. John A Lees, Simon R Harris, Gerry Tonkin-Hill, Rebecca A Gladstone, Stephanie W Lo, Jeffrey N Weiser, Jukka Corander, Stephen D Bentley, and Nicholas J Croucher. Fast and flexible bacterial genomic epidemiology with PopPUNK. Genome Res., 29(2):304–316, February 2019.

